# Ion-dependent structure, dynamics, and allosteric coupling in a non-selective cation channel

**DOI:** 10.1101/2021.06.16.448691

**Authors:** Adam Lewis, Vilius Kurauskas, Marco Tonelli, Katherine Henzler-Wildman

## Abstract

The selectivity filter (SF) determines which ions are efficiently conducted through ion channel pores. NaK is a non-selective cation channel that conducts Na^+^ and K^+^ with equal efficiency. Crystal structures of NaK suggested a rigid SF structure, but later solid-state NMR and MD simulations questioned this interpretation. Here, we use solution NMR to characterize how bound Na^+^ vs. K^+^ affects NaK SF structure and dynamics. We find that the extracellular end of the SF is flexible on the ps-ns timescale regardless of bound ion. On a slower timescale, we observe a structural change between the Na^+^ and K^+^-bound states, accompanied by increased structural heterogeneity in Na^+^. We also show direct evidence that the SF structure is communicated to the pore via I88 on the M2 helix. These results support a dynamic SF with multiple conformations involved in non-selective conduction. Our data also demonstrate allosteric coupling between the SF and pore-lining helices in a non-selective cation channel that is analogous to the allosteric coupling previously demonstrated for K^+^-selective channels, supporting the generality of this model.

## INTRODUCTION

Bacterial ion channels are convenient model systems for understanding the structure, dynamics and function of the pore domain of more complex eukaryotic channels. KcsA and MthK are two K^+^-selective, bacterial channels with homotetrameric pore structures. Each monomer is composed of two transmembrane helices, one pore helix, and the selectivity filter (SF), a structured loop with a highly conserved sequence that is responsible for selective K^+^ conduction.^1–3^ This general architecture is conserved in eukaryotic voltage-gated K^+^ channels.^4,5^ While much has been learned about conserved aspects of channel function from KcsA and MthK, these channels are K^+^-selective, potentially limiting the insight to be gleaned for non-selective channels such as HCN, CNG, or TRP channels, which differ from K^+^ channels in SF structure.^6,7^ NaK, on the other hand, is a bacterial channel that non-selectively conducts both K^+^ and Na^+^. The crystal structure of the NaK pore is very similar to KcsA and MthK (Fig. 1), differing only in the SF, where two of the four canonical ion binding sites are abolished.^8^ The biochemical tractability of systems like KcsA or NaK permits detailed biophysical studies aimed at understanding the fundamental and highly conserved processes of ion selectivity, permeation and channel gating. In the present study, we employ NaK as a model non-selective channel to ask how the structure and dynamics of the SF are affected by ion identity and to what extent ion-dependent conformational changes in the SF are propagated throughout the pore.

**Fig. 1.**
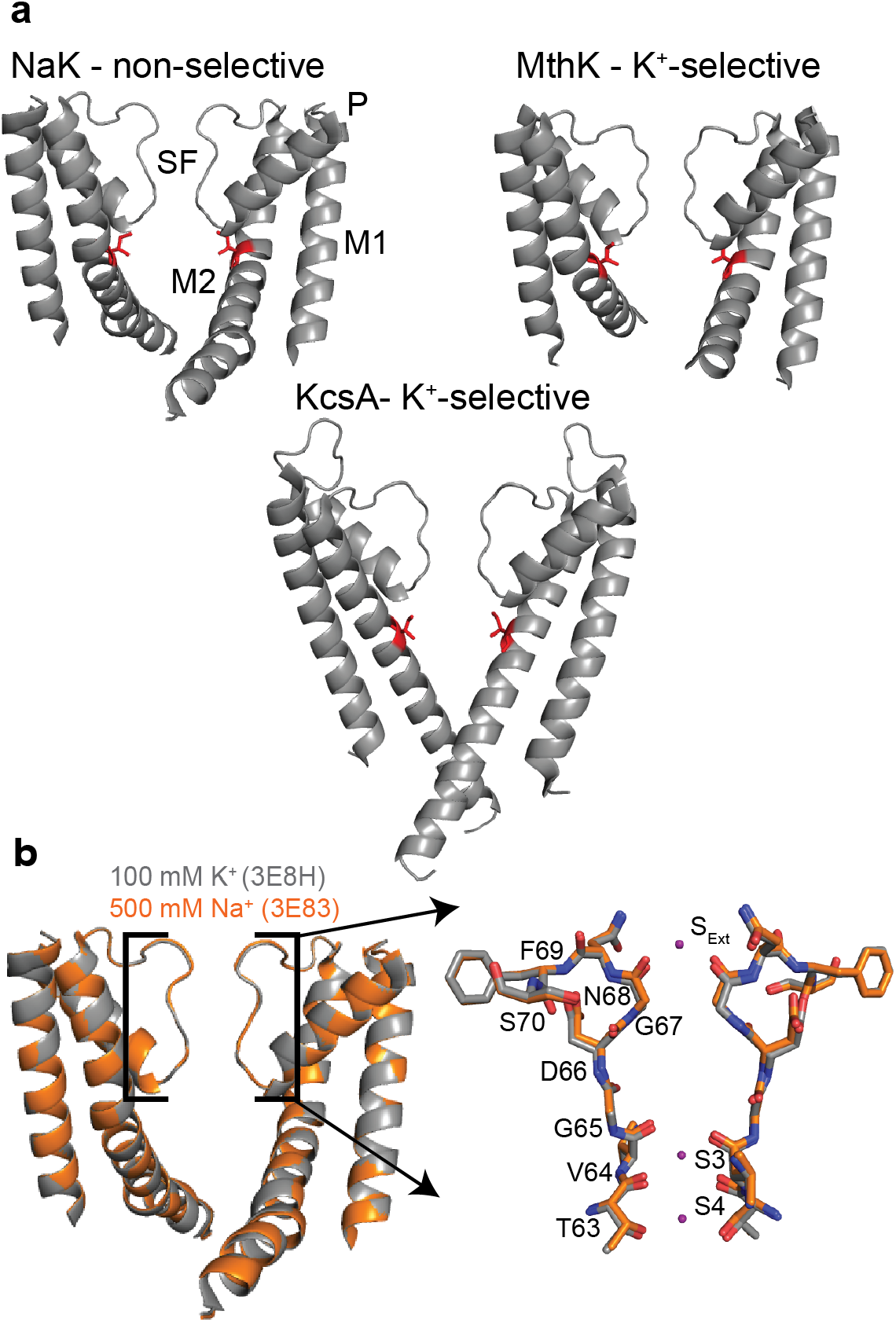
NaK channel structural comparisons. **a** Structures and selectivities of 3 of bacterial ion channels, NaK (3E8H), MthK (3LDC), and KcsA(1K4C). The selectivity filter (SF) along with M1, P, and M2 helices are labeled for NaK. Residue I88, I84,and I100 is highlighted in red on NaK, MthK, and KcsA structures respectively. **b** Overlay of NaK structures determined in 100 mM K^+^ (3E8H) and 500 mM Na^+^ (3E83). The selectivity filter (SF) along with M1, P, and M2 helices are labeled.The SF is enlarged on the right with residue labels. Ion binding sites are labeled and K^+^ ions modeled in the 3E8H structure are shown as well. In both panels, two opposing of four total subunits are shown for clarity.

Initial X-ray crystallographic studies of NaK suggested that the NaK SF conformation was rigid, as the SF structure was virtually unchanged when crystallized in 100 mM K^+^ or 500 mM Na^+^ (Fig. 1).^8,9^ This observation led to the hypothesis that K^+^ and Na^+^ bind in different modes to the same SF structure. The limited number of ion binding sites in the non-selective SF reduces the frequency of knock-on ion interactions proposed to drive K^+^-selective conduction in 4-ion-binding site SF structures^10,11^, thereby leading to non-selective conductance in NaK. However, MD simulations showed that the conformation of the NaK SF determined in 100 mM K^+^ (3E8H) was not conductive to Na^+^, suggesting that the SF must change conformation to conduct Na^+^.^12^ A combined ssNMR and MD study^13^ presented evidence that multiple SF states may be involved in Na^+^ permeation in NaK. SF residues V64 and G65 (Fig. 1) displayed ion-dependent chemical shifts, which was unexpected based on the identical conformation of these residues in Na^+^- and K^+^- bound crystal structures (Fig. 1). Na^+^ permeation was also not observed in MD simulations using the structure of NaK solved in 500 mM Na^+^ (3E83), and only occurred when the SF was rendered asymmetric by flipping the T62 carbonyl orientation in a subset of subunits. These lines of evidence suggest that the NaK SF is not static, and that structural heterogeneity in the SF on a relatively slow timescale may be important for achieving non-selective conductance of Na^+^ in addition to K^+^.

The structure of the SF in certain channels is also affected by allosteric coupling to the inner gate, and vice versa. Perhaps the most extensively characterized example is that of C-type inactivation in KcsA, where prolonged opening of the inner gate causes the SF to enter a ‘collapsed’ state that is non-conductive.^14–18^ Recently, it has also been proposed that SF-inner gate coupling controls channel activation for MthK and KcsA.^19,20^ In this model, the SF is in a non-conductive conformation when the inner gate opening is small, but upon further opening the SF widens to allow permeation. Residues on M2 at the base of the SF have been identified as mediators inter-gate coupling, specifically I100 and F103 for KcsA and I84 in MthK (analogous to I100 in KcsA). Upon prolonged opening, the KcsA SF undergoes a conformational change that results in inactivation. MthK has also been reported to inactivate, although the inactivation phenotype differs in MthK and KcsA.^17,20–23^

Here we experimentally determine how the NaK SF structure changes to accommodate Na^+^ vs. K^+^ and how SF dynamics are affected by the permeant ion on both fast (ps-ns) and slow (ms-s) timescales. As mentioned above, ssNMR data indicates V64 and G65 adopt ion-dependent structural states, but resonances were not observed for residues 66-70^13^ so it remains unclear how this region of the SF is affected by the permeant ion. We also assess whether the coupling mechanism observed in MthK and KcsA is also observed in NaK. Previously, our group has shown that NaK and NaK2K, a K^+^-selective mutant of NaK with a different SF structure^24^, have distinct NMR chemical shifts for residues far from the sites of mutation in the SF.^25^ This demonstrates that in principle the state of the putative inner gate can be influenced by the structure of the SF. Indeed, initial MD data presented for NaK2K suggests that the SF and M2 helix are coupled in a similar manner to KcsA or MthK.^20^ In ssNMR spectra of NaK, however, peaks for residues below V91 on M2 were not visible^13^, making it impossible to assess how or if the SF structure is communicated throughout the pore in the wild type channel.

Here we use solution NMR to investigate the ion dependence (Na^+^ vs K^+^) of SF structure, dynamics, and coupling to the inner gate in NaK solubilized in isotropic bicelles (q=0.33 DMPC/DHPC). For these studies we used a NaK construct lacking the first 18 residues. A very similar construct lacking the first 19 residues (NaKΔ19) has been used extensively in studies of ion selectivity in NaK^8,9,24,26,27^ (referred to as NaK throughout for clarity). Our solution NMR-based approach allows us to observe resonances throughout the NaK structure and characterize dynamics in a quantitative manner. Using ^15^N R_1_, R_1_ρ, and {^1^H}-^15^N NOE experiments, we show that residues 66-70 in the SF are highly dynamic on the ps-ns timescale regardless of bound ion. On a slower timescale, we observe chemical shift perturbations (CSPs) throughout the SF indicative of an ion-dependent conformational change that extends to the extracellular mouth. We also find evidence for increased structural heterogeneity in Na^+^. Several SF residues exhibit multiple resonances in Na^+^, while methyl-CPMG experiments showed that V64 experiences pronounced chemical exchange on the low ms timescale. Further, we observed CSPs far from the SF suggesting that the SF structure may be coupled to the rest of the pore. Methyl-methyl NOEs showed that the I88 side-chain, a residue critical for allosteric coupling in MthK and KcsA, adopts a different conformation in K^+^ vs. Na^+^. Together, these results indicate that the NaK SF can adopt ion-dependent structural states with similar dynamics on a fast timescale, but that ms-timescale exchange is increased in Na^+^. Further, the change in SF structure is communicated to the gating hinge, showing that allosteric coupling is intrinsic to NaK.

## RESULTS

### NaK SF flexibility differs markedly from the intracellular to extracellular end

To understand the impact of binding of Na^+^ and K^+^ on the structure and dynamics of the NaK channel, ^1^H-^15^N TROSY-HSQC spectra of U-^15^N,^2^H labeled NaK were collected in 100 mM K^+^ and 600 mM Na^+^. The high Na^+^ concentration was chosen to saturate Na^+^-binding to NaK in order to drive the SF towards a Na^+^-preferred state, and is based on previously reported Na^+^ binding constants determined by ITC^26^. Quantitative NMR studies of a protein the size of NaK solubilized in a bicelle require perdeuteration. In both ionic conditions, backbone amide resonances for residues 33-40, 53-65, and 79-87 are either weak or not visible due to lack of back-exchange of amide deuterons to protons after expression in D_2_O (Fig. S1). In order to monitor amides throughout NaK, we developed an SDS-based unfolding/refolding protocol to enhance back-exchange of amide protons (See Methods, Supplementary note 1). Spectra of refolded NaK solubilized in isotropic bicelles overlay very well with spectra of NaK that did not undergo the refolding protocol, but the peak intensity is increased for many weak peaks in the K^+^ sample and many more peaks are visible in the Na^+^ sample (Fig. S1). Thus, our SDS back-exchange method did not alter the conformation of NaK, but does allow us to observe and assign nearly all backbone amides, enabling detailed investigation of NaK structure and dynamics.

With more backbone amides in the SF visible in the NMR spectrum as a result of our new purification scheme, we noticed that SF residues V64 and G65 were poorly back-exchanged in both ionic conditions, suggesting that the amides for these residues are not water-accessible. Conversely, intense resonances were observed for residues 66-70 at the mouth of the SF, leading us to hypothesize that the dynamic character of the SF differs markedly along the intracellular to extracellular axis. We performed ^15^N R_1_, R_1_ρ, and {^1^H}-^15^N NOE measurements^28^ using refolded samples in both 100 mM K^+^ and 600 mM Na^+^ to more quantitatively characterize ps-ns timescale internal dynamics in NaK (Supplementary note 2, Fig S2). R_1_ and NOE data are shown in Fig. 2. For folded proteins with little internal motion, these quantities are consistent for residues found in stable secondary structure, with the precise value depending on the global tumbling of the protein molecule (or in our case, protein plus bicelle). Indeed, measured R_1_, R_2_, and NOE values are quite uniform for residues predicted to be in α-helices based on the crystal structure, suggesting that NaK remains well-ordered in our NMR samples (Fig 2, Fig. S2). Deviations from the average value for helical residues are indicative of the presence of internal dynamics, with increased R_1_ values and decreased NOE values indicating enhanced dynamics.^28,29^

**Fig. 2.**
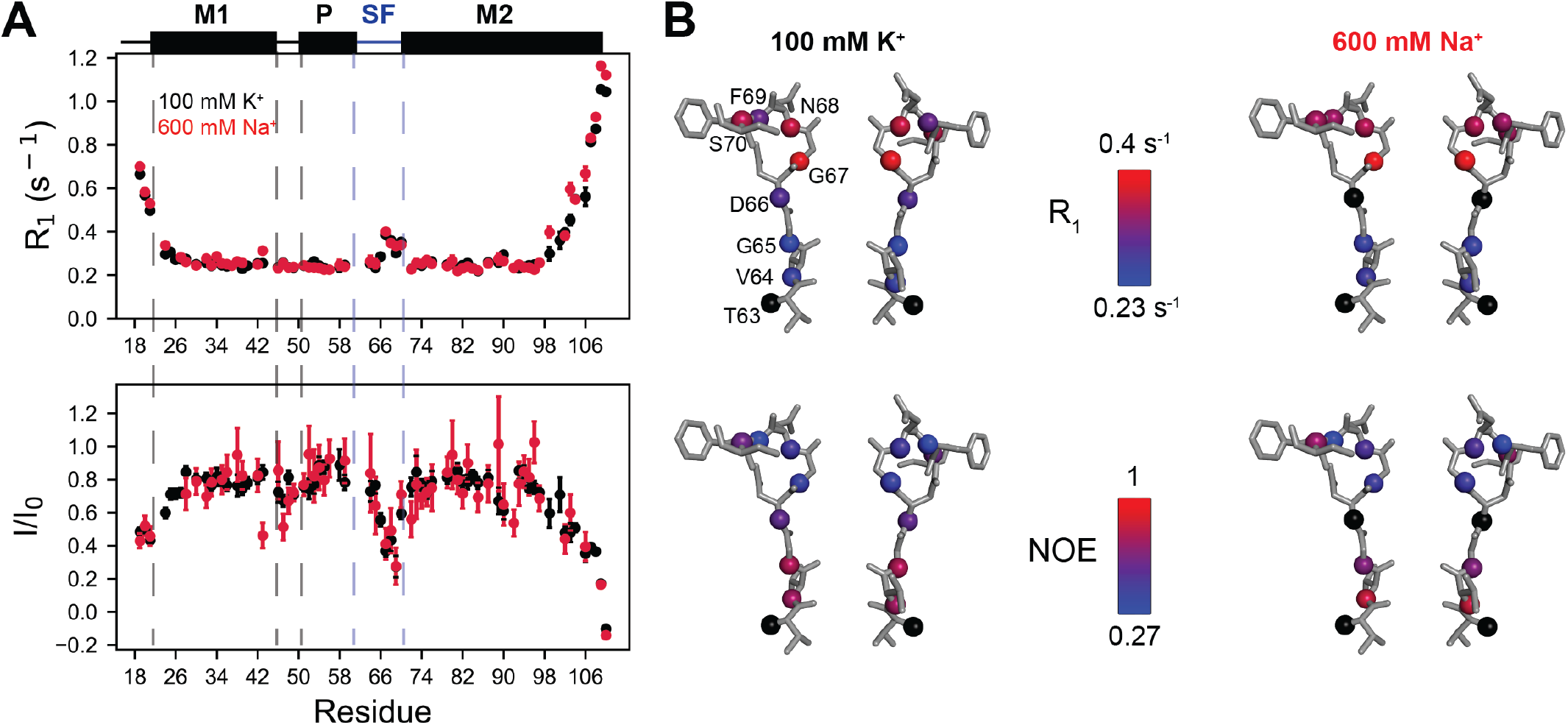
Gradient in ps-ns dynamics across NaK SF. **a** Backbone amide ^15^N R_1_ and {^1^H}-^15^N NOE values recorded at 750 MHz for NaK in 100 mM K^+^ (black) and 600 mM Na^+^ (red). Secondary structure elements are shown above, where helices are rectangles and loops are lines. The boundaries between elements are based on the 3E8H crystal structure. Residues 63-70 are indicated by the blue lines and are shown as sticks in B. **b** Relaxation data plotted on the NaK SF (showing 2 of 4 opposing subunits). Backbone N atoms are shown as spheres and are colored according to the scale indicated for each parameter. Residues for which data are unavailable due to resonance overlap or lack of assignment are colored black.

It is clear that in both K^+^ and Na^+^ there is a pronounced gradient in ps-ns dynamics from the intracellular to extracellular end of the SF. Parameter values for V64 and G65 are similar to those for residues found in the transmembrane helices. V64 contributes to the canonical S3 and S4 binding sites, while G65 is adjacent to S3, and based on our relaxation data it is likely that these sites are stably maintained in solution. Conversely, there is a pronounced drop in the NOE and increase in R_1_ from residues 66-70, indicating that the backbone in this region is more flexible than at the intracellular end of the SF. In particular, residues 67-69 have very elevated R_1_ values and markedly lower NOE values indicative of extensive ps-ns timescale dynamics at these sites. This data is not consistent with the interpretation of a rigid SF structure, as suggested based on X-ray crystallography.^9^ Not only is the NaK SF dynamic on a slow timescale, as shown previously by ssNMR^13^, it is also dynamic on a fast timescale. This conclusion is consistent with our H/D exchange data showing that residues 66-70 more readily exchange with bulk water (Fig. S1) and with the fact that correlations were not observed for residues 66-70 in dipolar-coupling based ssNMR experiments where structural heterogeneity impedes magnetization transfer.^13,30^ Thus, our data clearly demonstrates that the extracellular mouth of the SF is structurally labile, potentially enabling this region to adapt to different types of bound ions.

### The SF conformation is ion-dependent

Chemical shifts measured for NaK in Na^+^ and K^+^ were compared to further understand how ion identity affects the SF structure on a slower (ms-s) timescale. Backbone amide ^1^H and ^15^N chemical shifts were obtained from 2D TROSY-HSQC spectra while C’ and Cα chemical shifts were obtained from TROSY-HNCO and HNCA spectra respectively. Additionally, ILV methyl chemical shifts were obtained from ^1^H-^13^C HMQC spectra of U-[^2^H,^15^N] Ile(δ1)-^13^CH_3_, Leu/Val-(^13^CH_3_/^12^C^2^H_3_) labeled NaK. Overlays of 2D TROSY-HSQC and ^1^H-^13^C HMQC spectra obtained in Na^+^ and K^+^ are shown in Fig. S3.

Residues 64-70 in the SF region experience significant amide chemical shift perturbations (CSPs) between 100 mM K^+^ and 600 mM Na^+^ (Fig. 3). Significant perturbations to Cα chemical shifts also occur for resides 66 and 68-70 while all residues in this region other than N68 exhibit perturbations to C’ greater than 0.2 ppm. Differences in chemical shifts between K^+^ and Na^+^ could in principle result from changes in ion occupancy/coordination for residues in close proximity to the ion binding sites, or from a structural change in this region. While carbonyl oxygen atoms are the ion-coordinating ligands in the SF, of the residues 66-70 only G67 C’ is observed to bind ions in the crystal structures, at a site termed S_ext_ at the extracellular mouth (Fig. 1).^8,9^ The backbone carbonyls of D66 and N68 have been reported to bind ions at a site S_side_ in MD simulations.^13^ A density assigned to a Na^+^ ion was also resolved at this site in another recently published crystal structure.^31^ The extent of backbone CSPs in the SF beyond these residues are difficult to explain solely based on changes in ion occupancy. Additionally, CSPs are observed for the methyl groups of V64, V59, L48, and I84, located behind the SF. Notably, the methyl groups of V64, V59, and I84 are not expected to be water (or ion) accessible based on the crystal structure.^8,9^ Together, all of these chemical shift perturbations support the existence of an ion-dependent conformational change in the SF, with residues 65-70 adopting distinct structural states in 100 mM K^+^ and 600 mM Na^+^.

**Fig. 3.**
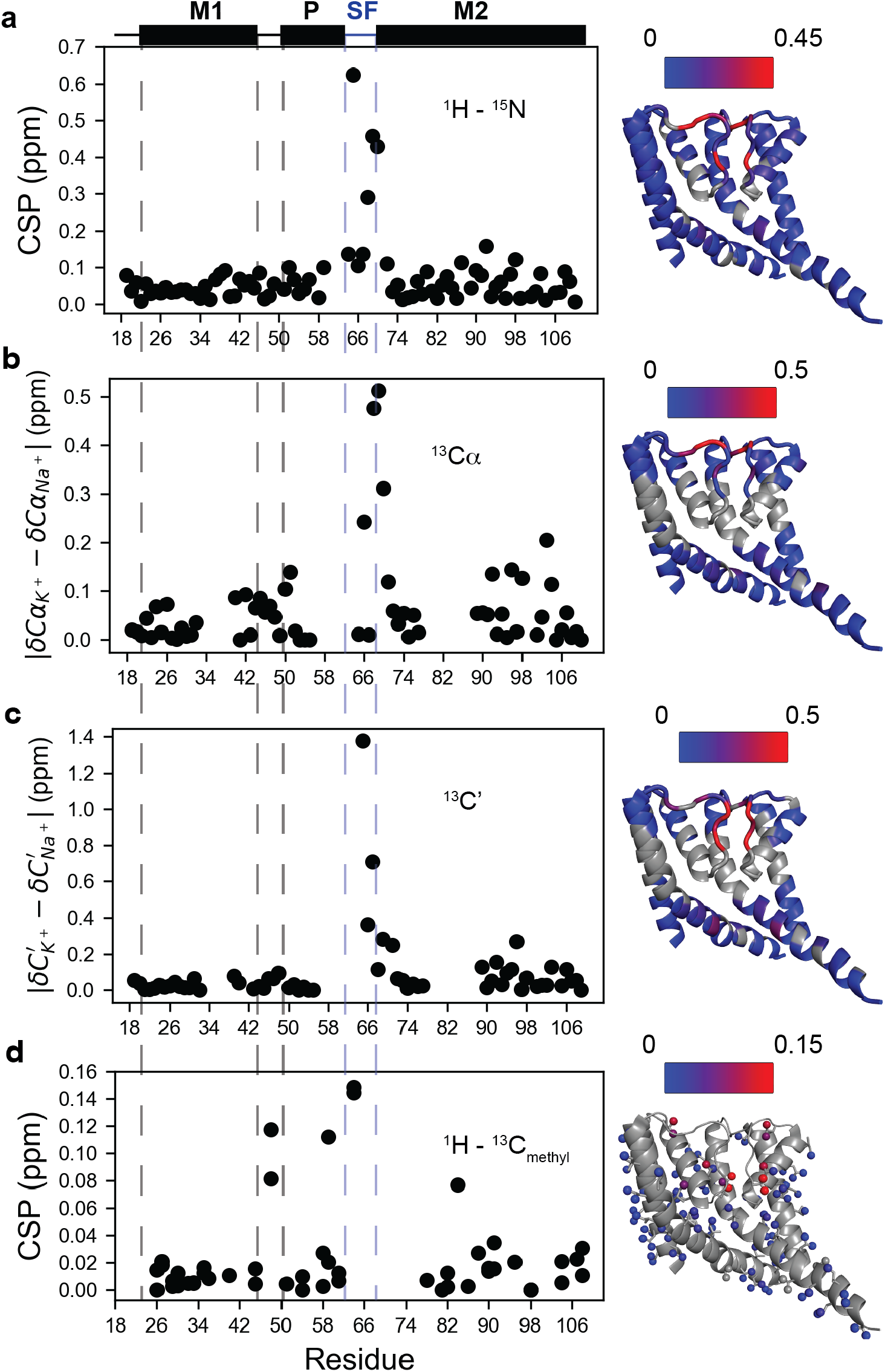
SF and hinge residues display ion-dependent chemical shifts. Backbone amide (**a**) and ILV methyl (**d**) CSPs along with ^13^C’ (**c**) and ^13^Cα (**b**) chemical shift differences were calculated for NaK in 100 mM K^+^ vs 600 mM Na^+^. CSPs (calculated as described in Methods) and Δδ values for residues assigned in both conditions are plotted against residue number and on the NaK structure (3E8H, 2 adjacent of 4 subunits shown for clarity). Scale bars are indicated for each plot. ^1^H-^15^N CSPs, ^13^C’ Δδ, and ^13^Cα Δδ values are plotted on the backbone and residues for which data are not available are colored gray. ^1^H-^13^C_methyl_ CSPs are plotted on the side-chains and the backbone is shown in gray.

Furthermore, several residues in the SF region of NaK were observed to have multiple resonances. More residues display additional resonances in Na^+^ than in K^+^, indicating a greater number of states in Na^+^-saturated NaK. No peak doubling is observed in amide spectra in K^+^, but spectra collected in Na^+^ display minor resonances for G67, F69, and S70. These three residues undergo large CSPs upon changing ionic conditions (Fig. 4). For methyl resonances, the γ1 and γ2 methyl groups of V64 and V59 are each split into two partially overlapped resonances (Fig. 4) in Na^+^, while only having a single resonance each in K^+^. The δ methyl groups of L48 display at least two resonances in both ionic conditions. Interestingly, the major state for L48b is inverted between K^+^ and Na^+^. In K^+^, the major state occurs at ∼0.3 ppm in ^1^H and has a population of ∼86% while the minor peaks around 0.4 and 0.2 ppm have populations of ∼7% each. In Na^+^, the major state occurs ∼0.2 ppm, rather than 0.3, and has a population of ∼82%. Only a single minor state is observed in this case. L48a displays three resonances in both ionic conditions. In K^+^, again, the major state makes up 86% of the population while each minor state is ∼7%; however, the populations in Na^+^ are more evenly distributed with population of 47%, 37%, and 16%.

**Fig. 4.**
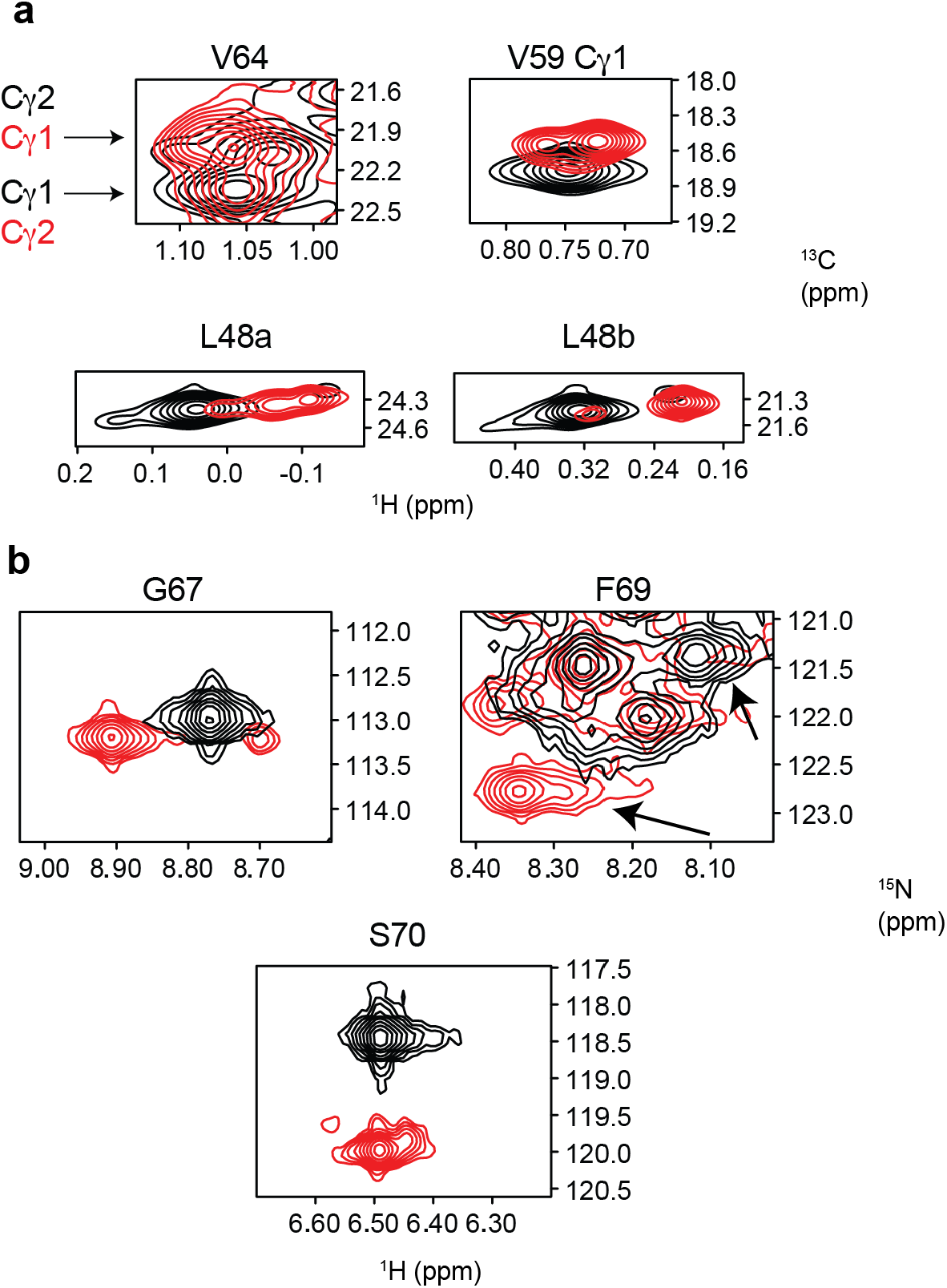
Na^+^ increases heterogeneity in the SF. Overlays of SF peaks from (**a**) ^1^H-^13^C HMQC and (**b**) TROSY-HSQC spectra of NaK in 600 mM Na^+^ (red) and 100 mM K^+^ (black). Stereospecific assignments based on NOESY data are given where possible.

Collectively, these data indicate that SF adopts distinct states in K^+^ and Na^+^. The backbone CSP data suggest a structural change occurring in the SF at the extracellular mouth between residues 65-70, and methyl CSPs for residues 64, 59, and 84 suggest that the change in structure is propagated in the vicinity of the SF. The data also indicate that several minor SF states exist in Na^+^. The CSPs we observe in the vicinity of the SF upon switching ion are largely consistent with very recent ssNMR data reported for NaK in liposomes^30^, further emphasizing that the structure and dynamics of NaK are essentially the same in bicelles and liposomes. To more precisely and quantitatively characterize the ion-dependent changes in SF structure and slow-timescale dynamics in the SF region and how the impact of ion binding propagates further down the NaK pore-lining helices, we turned to methyl NOESY and CPMG experiments.

### The M2 helix is coupled to the SF

Structural coupling between the SF and the inner gate is frequently observed in K^+^-selective channels and structurally related channels, such as NaK. Do the structural changes in the SF we detect in Na^+^ versus K^+^ propagate through the M2 helix? The first piece of evidence for structural coupling is that, while the largest CSPs occur near the SF, there are more subtle but still significant perturbations that occur for residues 87-94 around the M2 hinge (Fig. 3) between the Na^+^ and K^+^ bound states. ^1^H-^15^N spectra CSPs are evident for G87 and L90-F92. Backbone carbon CSPs for L90 and V91 are fairly small, but more pronounced shifts occur for I88, G89, and F92. CSPs are also observed for the side chain methyl groups of I84, I88, and V91 (Fig. 3, Fig. S3). Several residues near the bundle crossing proposed to be the cytoplasmic gate of NaK^32,33^ also exhibit chemical shift differences between Na^+^ and K^+^, including the amides for L98 and Q103 and a methyl resonance ambiguously assigned to either V100 or V102.

Methyl-methyl NOESY data was collected on ILV labeled NaK samples in K^+^ and Na^+^ to characterize these potential structural changes in more detail. In general, the NOEs for both samples are consistent with the overall architecture of NaKΔ19 observed in crystal structures. However, it is clear that the side chain of I88 adopts a different conformation in K^+^ vs. Na^+^ (Fig. 5). This is interesting because I88 is located in the M2 helix in a position analogous to I100 in KcsA and I84 in MthK, two residues that have been implicated in inter-gate allosteric coupling.^17,20–22^ The I88 δ1 NOEs to V64 γ2 and I84 δ1 are much stronger in K^+^ than Na^+^. In addition, I88 δ1 has a stronger NOE to V91 γ1 than V64 γ2 in Na^+^, while the relative intensities of these two NOEs are reversed in K^+^. In crystal structures of NaK, the shortest predicted intermethyl distance for I88 δ1 is to I95, consistent with our observation of a strong NOE to I95 under both conditions. In the structure of NaK in 100 mM K^+^ (3E8H), the I88 side chain is modeled in two conformations, one with Cδ pointing up and one with Cδ pointing down; whereas in the 3E83 structure of NaK in 500 mM Na^+^, I88 is only modeled in the down conformation. In the down conformation, I88 δ1 is closer to V91 γ1 than V64 γ2, consistent with the observed NOE intensities in Na^+^. On the other hand, I88 δ1 in the up conformation is closer to V64 γ2 than V91 γ1. Our data are therefore consistent with I88 adopting the down conformation in Na^+^ and the up conformation in K^+^.

**Fig. 5.**
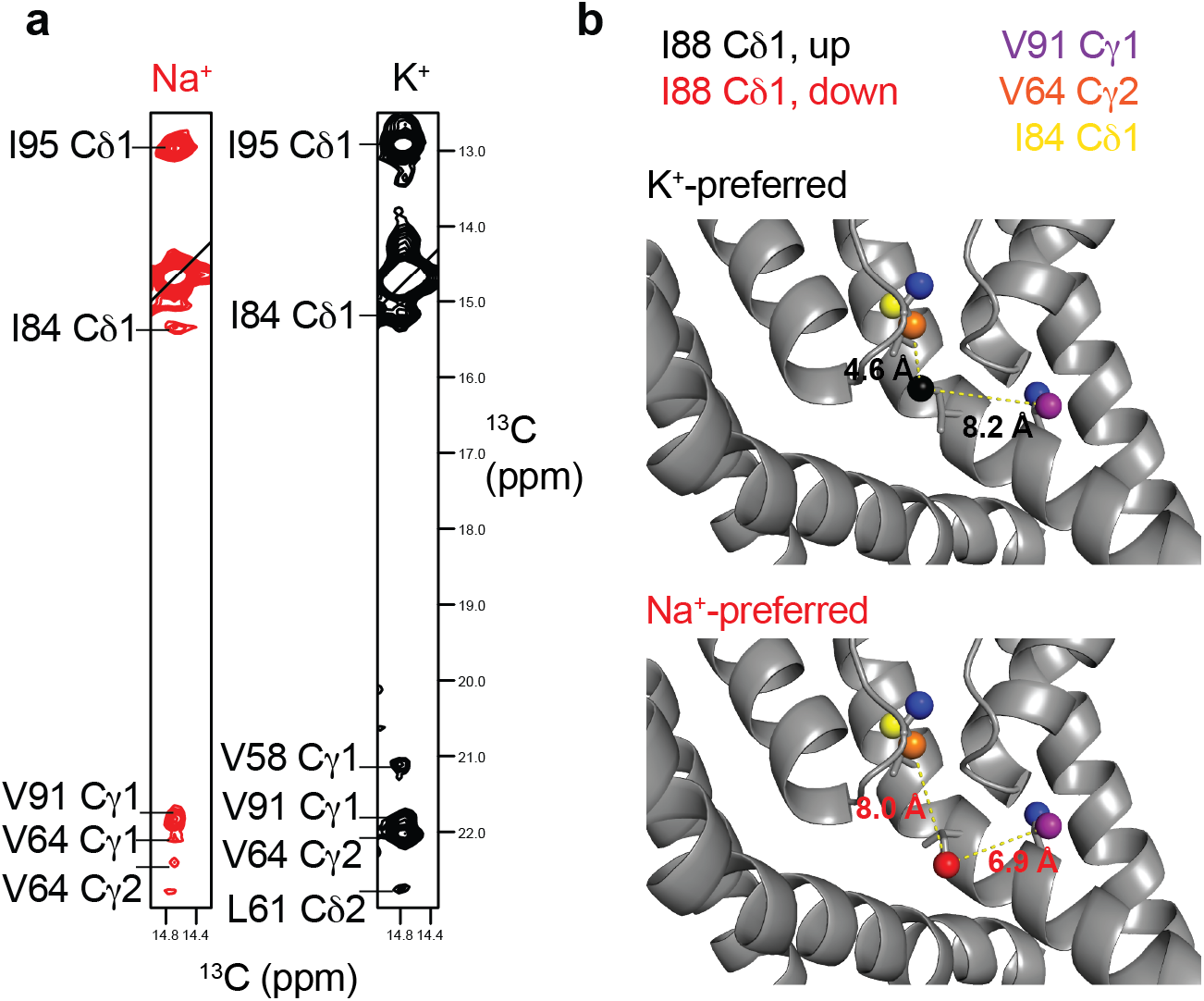
I88 side chain conformation is ion dependent. **a** Strips for I88 Cδ1 from 3D CCH NOESY experiments collected in 600 mM Na^+^ (red, ^1^H ppm = 0.567) and 100 mM K^+^ (black, ^1^H ppm = 0.576). **b** Distances from I88 Cδ1 to V64 Cγ2 and V91 Cγ1 in the ‘up’(black) and ‘down’(red) conformations. NOESY data collected in K^+^ is consistent with the ‘up’ conformation, while the data collected in Na^+^ is more consistent with the ‘down’ conformation.

Recently reported ssNMR data supported ion-dependent structural changes in NaK solely in the vicinity of the SF.^30^ In ssNMR spectra of NaK, resonances are not observed for residues 91-110, making it impossible to conclude how this region is affected by the permeant ion. Because we can resolve resonances throughout the NaK structure, we see that residues in the M2 helix show CSPs indicative of a structural change between the Na^+^- and K^+^-bound states. Additionally, we are not limited to observing changes in backbone structure because of our methyl-labeling approach. Indeed, we can see that methyl resonances for V59 in the P helix and I84, I88, and V91 in M2 are affected by bound ion identity and methyl NOESY spectra reveal that the I88 side-chain conformation switches between Na^+^ and K^+^.

### Na^+^ enhances ms timescale dynamics in the SF

We noticed broadening of several NaK methyl resonances, including I84, I88, and I95 on M2, as well as V64 in the SF, both in the presence of K^+^ and Na^+^. Such line broadening can be indicative of ms-timescale conformational exchange, hence ^13^C Multiple-Quantum (MQ) CPMG experiments were performed to further characterize these potential dynamics.^34^ In K^+^, several methyl groups had significant exchange contributions to the MQ relaxation rates (R_ex_) (> 3 Hz) (Fig. 6). Interestingly, most of these residues are localized to the hinge and base of the SF, including I88, L90, and V91 on M2 as well as L35 on M1. V58 in the P helix and V64 Cγ2 in the SF also show R_ex_ >3 Hz. More quantitative analysis of exchange rates and populations was hindered by the relatively small size of the exchange contribution for these residues (~4-5 Hz at 800 MHz). However, the fact that the majority of residues with significant R_ex_ contributions are found at the hinge and base of the SF further supports the notion that coupling between the SF and M2 is inherent to NaK and is mediated by the SF-hinge interface.

**Fig. 6.**
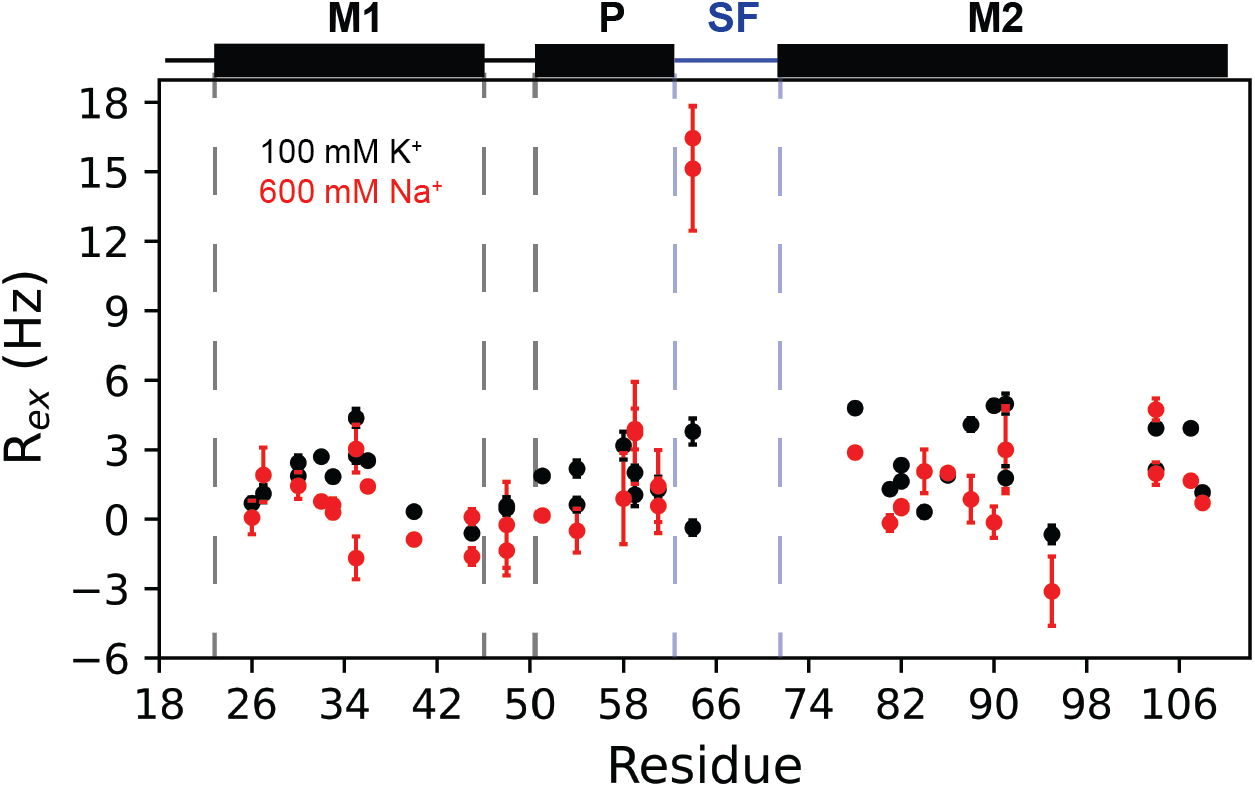
V64 undergoes enhanced chemical exchange in the presence of Na^+^. R_ex_ values for ILV labeled NaK in the presence of Na^+^ and K^+^ are plotted against the sequence. Note the marked incrased in R_ex_ for V64 in the presence of Na^+^.

Surprisingly, CPMG experiments collected for NaK in the presence of Na^+^ revealed a marked increase in R_ex_ for V64 Cγ1 (none to ∼15Hz) and Cγ2 (~4 to ∼16 Hz), as well as V59 Cγ2 (∼2 to ∼4Hz) (Fig. 6). Other hinge residues display slightly lower R_ex_ in Na^+^ than K^+^; however, these small differences may not be significant due to the lower S/N in the Na^+^ dataset. Due to the importance of V64 in ion coordination, a second ^13^C MQ CPMG data was collected at 600 MHz along with ^1^H Single Quantum (SQ) CPMG datasets at 900 and 600 MHz for the Na^+^ sample to attempt to quantify the exchange rate.^34,35^ Unfortunately, interfering signals from residually protonated DMPC present in the bicelle prevented accurate extraction of V64 peak intensities in some spectra. The ^1^H SQ CPMG experiment did, however, reveal significant ^1^H R_ex_ for L48 Hδ atoms that point behind the SF. Global fitting was performed for V64 and V59 ^13^C MQ and L48 ^1^H SQ data to extract an estimate for the rate constant for chemical exchange in the SF, k_ex_ (assuming two state exchange). Satisfactory fits could not be obtained when V64, V59, and L48 were fit globally, likely reflecting multistate exchange as suggested by the three resolved amide resonances for some residues in this region. However, pairwise fitting of V64 and V59 MQ data or V59 MQ and L48 SQ was successful, yielding k_ex_ = 390 ± 47 s^-1^ and k_ex_ = 356 ± 67 s^-1^, respectively (Fig. S4). Due to the small number of methyl groups and lack of clear evidence to support two-state exchange, we did not attempt to extract populations. In summary, residues in the SF, especially ion-coordinating residue V64, are seen to undergo enhanced chemical exchange in the presence of 600 mM Na^+^ as opposed to 100 mM K^+^, with a timescale of approximately 350 s^-1^ in Na^+^.

## DISCUSSION

There is mounting evidence suggesting that multiple SF conformations are needed for non-selective conduction in certain ion channels.^13,36^ Additionally, while SF-inner gate coupling has been demonstrated for some K+ channels^17,20,21^, questions remain regarding how general this type of coupling may be across other channel families. In the present study, we utilized the non-selective, prokaryotic ion channel NaK to characterize how the structure and dynamics of the SF respond to the identity of the ion occupying the SF and how any changes are allosterically communicated to other parts of the channel.

We found that the NaK SF is structurally labile and that the SF conformation is ion-dependent (Fig. 3). This notion is in contrast to crystal structures of NaK where the SF structure is identical in 500 mM Na^+^ and 100 mM K^+^ and in agreement with a prior ssNMR study of NaK where V64 and G65 had ion dependent chemical shifts (Fig. 1).^13,30,37^ Our observation of enhanced ps-ns dynamics for D66-S70 also explains the difficulty observing this region in ssNMR spectra. In line with the backbone flexibility, we observe extensive backbone CSPs in this region between K^+^ and Na^+^-bound samples. Observed CSPs are in accord with two recent structures of NaK solubilized in DM – a microED structure solved in Na^+^ (6CPV)^38^ and a new X-ray structure solved in mixed ion conditions (6V8Y).^31^ In both structures, N68 is seen to adopt alternative conformations that result in changes to backbone ϕ and ψ angles for N68 and F69, as well as slight reorientation of the F69 sidechain. Accordingly, we see large CSPs for N68 HN, Cα, and C’ and F69 HN, Cα, and C’ between Na^+^ and K^+^ (Fig. 3). In fact, the perturbations to Cα for N68 and F69 are the two largest for all Cα’s from residues 65-70 in our samples. The new X-ray structure also shows a reorientation of the carbonyl group for D66 in one protomer.^31^ This reorientation largely affects the conformation of D66 C’ and G67 HN. Again, we see large perturbations to both D66 C’ and G67 amide chemical shifts. Thus, our CSP data collected in a lipidic environment are consistent with the perturbations to the canonical SF structure observed in detergent-solubilized structures of NaK.

The dynamic nature of the NaK SF has important implications for the permeation and non-selectivity mechanisms of NaK and other non-selective ion channels. Our relaxation data are in accord with MD simulations of the non-selective channels NaK, NaK2CNG, and TRPV1 showing that these SFs are highly dynamic.^13,36,39–41^ Interestingly, the TRPV1 SF was reported to adopt different structural states during K^+^ and Na^+^ permeation.^36^ The activated state structure of the TRPV1 SF (3J5Q) was stable during K^+^ permeation, but an asymmetric deformation occurred during Na^+^ permeation as a result of Na^+^ binding to an aspartate adjacent to the conduction path during the simulation. Interestingly, a putative Na^+^ binding site behind the SF was proposed to stabilize the alternate conformation of D66 seen in the recently published 6V8Y structure of NaK.^31^ The break in SF symmetry in TRPV1 is also is reminiscent of results for NaK, where Na^+^ permeation only occurred in forced asymmetric states^13^, although Na^+^ could permeate the TRPV1 SF in both symmetric and asymmetric conformations.^36^ The appearance of multiple states for several SF residues in the presence of Na^+^ could indicate a break in SF symmetry in our NMR samples and support the notion that Na^+^ may permeate NaK in an asymmetric filter conformation.

Considering these results for TRPV1 and NaK, it appears that conformational flexibility in the SF is important for non-selective conduction, especially in the case of the Na^+^. SF plasticity may allow both K^+^ and Na^+^ to permeate efficiently in different conformations. For example, SF flexibility may allow Na^+^ to permeate in a hydrated or partially hydrated state, as suggested by simulations of NaK2CNG-N.^11^ Because dehydration is more unfavorable for K^+^ than Na^+^, allowing Na^+^ to permeate while hydrated may contribute to non-selective conductance. This notion is in accord with different MD simulation studies showing that, in general, ion selectivity is decreased as ion hydration in the SF increases.^11,41^ On the other hand, recent structures of human CNGA1 solved in a range of ionic conditions showed no significant structural change in the SF.^42^ It is possible that the reported structures do not capture all states necessary for permeation, as was the case with NaK. Alternatively, it may be that since the CNGA1 SF is wider than in NaK, hydrated ions can permeate without any structural change. A similar hypothesis has been proposed for several TRP channels that display wide SF openings even in closed states of the channel.^7^ Other non-selective channels with an available structure, such as TAX-4^43,44^ or HCN1^45^, were solved in a single ionic condition, limiting potential insight into how the SF adapts to different types of ions. Additional structures of non-selective channels in a variety of ionic conditions, as well as simulations of these structures, will be needed to come to a complete understanding of non-selective permeation. In any case, our data demonstrate significant dynamics on multiple timescales within the NaK SF, supporting the notion that for non-selective channels with a narrow pore, such as NaK, flexibility of the SF is required to allow conduction of both K^+^ and Na^+^.

We observed a Na^+^-promoted increase in ms-timescale conformational exchange for L48, V59, and V64, where the increase in R_ex_ for V64 was especially large (Fig. 6, S4). This result is intriguing given that the residue analogous to V64 in MthK^20,46^ (V60) and KcsA^15,18,47^ (V76) has been reported to flip such that the ion-coordinating carbonyl group points away from the conduction path. Moreover, the E71A mutant of KcsA crystallized in the flipped conformation in high Na^+^ and was less K^+^-selective than WT KcsA, suggesting that carbonyl flipping could be an important contributor to non-selectivity.^18^ A previous study of NaK proposed that carbonyl flipping occurs at residue T62, which could in principle result in exchange-induced line-broadening of V64; however, our NMR data cannot distinguish between these possibilities.^13^

We found that not only is the NaK SF structure ion dependent, but that the structural perturbation extends to M2 via I88 (Fig. 3, Fig. 5). This is significant because the analogous residue in MthK (I84) and KcsA (I100) has been implicated in mediating allosteric coupling between the SF and intracellular gate for these channels (Fig. 1).^17,20–22^ For MthK and KcsA, inter-gate coupling is implicated in both activation and inactivation.^15,16,19,20,23^ The activating stimulus for NaK is not currently known, nor is it known whether or not NaK undergoes inactivation in a manner similar to KcsA. Thus, we cannot say for certain which processes are influenced by SF-hinge coupling in NaK; however, our data suggests that I88-mediated allosteric coupling extends to NaK and supports a broader mechanism of allosteric regulation for channel pore domains.

In conclusion, we have shown that the NaK SF displays a large degree of structural plasticity on multiple timescales. We find the SF exhibits strong dynamics on the ps-ns timescale as well as the ability to adopt multiple ion-dependent states on a slower timescale. CSP data indicate that the slower, ion-dependent conformational change occurs between residues 65-70. These observations are in keeping with results from other non-selective channels showing that multiple SF conformations contribute to non-selective conduction. We also identify a residue located at the M2 hinge in NaK, I88, that is likely responsible for coupling the SF conformation to the inner gate. The analogous residue has been implicated in allosteric coupling in structurally related channels, suggesting that this mechanism of coupling may be quite general.

## METHODS

### Protein expression, purification, and reconstitution

A NaK construct lacking the first 18 residues was used for this study. This construct is very similar to NaKΔ19, which has been extensively characterized by X-ray crystallography and NMR previously.^25,33^ Growth and expression of isotopically labeled NaK was carried out as previously described.^25^ For ^15^N relaxation experiments, uniform ^15^N, ^2^H labeling was achieved through the use of M9 minimal media supplemented with 1 g/L ^15^NH_4_Cl, 4 g/L ^13^C, ^2^H glucose, and 0.5 g/L ^15^N, ^2^H isogro (Sigma-Aldrich) in D_2_O. The same isotopically labeled compounds were used to produce ^13^C-^1^H_3_ ILV labeled samples, but the amount of ^15^N, ^2^H isogro was reduced to 0.25 g/L and ∼140 mg/L 2-keto-3-(methyl-d_3_)-butyric acid-4-^13^C, 3-d and ∼70 mg/L 2-ketobutyric acid-4-^13^C,3,3-d_2_ were added ∼30 mins prior to induction.

An SDS-mediated refolding protocol was used for preparing ^15^N relaxation samples in Na^+^ and K^+^. This protocol was inspired by other membrane protein refolding studies, where refolding from the SDS-solubilized state has proven fruitful of a-helical membrane proteins.^48^ Cells were lysed by sonication in lysis buffer (100 mM NaCl, 200 mM KCl, 2.5 mM MgSO_4_, 20 mM Tris pH 7.5, DNase, 1 μg/mL pepstatin, 10 μM leupeptin, 100 μM PMSF) with 250 mM sucrose and 1 mg/mL lysozyme. The membrane fraction was isolated by centrifugation at 30,000 x g for 1 hour. The membrane fraction was then resuspended in lysis buffer + 0.5% SDS and allowed to rotate overnight to solubilize and to allow for back-exchange of deuterons to protons at exchangeable sites. Solubilized NaK was then purified via IMAC using Talon cobalt affinity resin (Clontech). For the K^+^ sample, the resin was equilibrated with Buffer A1 (300 mM NaCl, 5mM DM (decyl maltoside, Anatrace), 20 mM Tris pH 8). NaK was then incubated with the resin for 20-30 mins before collecting the flow through and washing with Buffer A1. The resin was then washed with Buffer A2 (100 mM NaCl, 200 mM KCl, 5mM DM, 20 mM Tris pH 8) and Buffer B (Buffer A2 + 5 mM imidazole). The resin was then incubated with Buffer C (Buffer A2 + 300 mM imidazole) for 20-30 mins and the elution was collected. For the Na^+^ sample, the same process was used, but the wash with Buffer A2 was replaced by an additional wash with Buffer A1 and Buffers B and C were prepared using Buffer A1. Elutions were concentrated to 0.5 mL and loaded on a Superdex 200 column pre-equilibrated with FPLC buffer. For the K^+^ sample, FPLC buffer consisted of 100 mM MOPS, 200 mM K^+^, 5 mM DM pH 7. For the Na^+^ sample, FPLC buffer consisted of 50 mM MOPS, 600 mM NaCl, 5 mM DM pH 7. Subsequently, FPLC fractions were evaluated using SDS-PAGE and selected fractions were pooled and incubated with thrombin to cleave the His-tag. Cleavage was monitored by SDS-PAGE.

Reconstitution into q = 0.33 DMPC/DHPC bicelles was performed using our previously published protocol with a few alterations.^25,49^ DMPC powder (1,2-dimyristoyl-sn-glycero-3-phosphocholine, Avanti Polar Lipids) hydrated with NMR buffer (for K^+^ - 100 mM MOPS, 100 mM K^+^, pH 7; for Na^+^ - 50 mM MOPS, 600 mM Na^+^, pH 7) at 20 mg/mL was bath sonicated for ∼1 min, incubated at ∼40ºC for 1 hour then bath sonicated again for 5 minutes. NMR buffer + 100 mM DM was then added to the DMPC to a final DM concentration of 10 mM and allowed to incubate for 15 mins, after which the lipid was incubated with the pooled NaK FPLC fractions for 30 mins. To remove detergent and form proteoliposomes, one aliquot of Amberlite XAD-2 (BioRad) was added and allowed to incubate for 1 hour, followed by addition of another aliquot and overnight incubation. The following day, a final aliquot was added and allowed to incubate for 1 hour. The total amount of Amberlite XAD-2 used to remove detergent was equal to 90 mg Amberlite per mg of DM in the sample. NaK proteoliposomes were then collected and pelleted by ultracentrifugation at 150,000 x g for 2 hours at 6ºC. The pellet was solubilized using DHPC (1,2-dihexanoyl-sn-glycero-3-phosphocholine, Avanti Polar Lipids) dissolved in NMR buffer at a molar ratio of 3:1 relative to DMPC. The sample was then subjected to four freeze-thaw cycles to produce homogenous bicelles. The Na^+^ sample was reconstituted using a molar ratio DMPC:NaK monomer of 95:1 while the K^+^ sample was reconstituted using a molar ratio DMPC:NaK monomer of 60:1.

Other NaK samples were prepared in a similar fashion; however, they were solubilized using lysis buffer + 20 mM DM rather than SDS. The samples were allowed to solubilize overnight before IMAC. For Na^+^ samples, IMAC was performed as described above. For K^+^, the initial wash with Buffer A1 was skipped, but was otherwise performed in the same manner. After IMAC, imidazole was removed using a PD10 desalting column (GE Healthcare) equilibrated with Buffer A1 for Na^+^ or Buffer A2 for K^+^. Samples were then incubated overnight with thrombin to cleave the His-tag. Samples were further purified by gel filtration chromatography as described above.

Reconstitution was performed nearly as described above for ^15^N, ^2^H labeled samples, but with slight alterations for methyl labeled samples. d54-DMPC (Avanti Polar Lipids) was used for methyl labeled samples, where preparation and proteoliposome formation was carried out as above. An aliquot of d22-DHPC (Avanti Polar Lipids) stored in chloroform was dried and then washed three times with pentane before lyophilizing overnight. The next day, the dried d22-DHPC was dissolved in either K^+^ or Na^+^ NMR buffer and used to solubilize proteoliposomes as described above. Lipid:protein ratios used for reconstitution of non-SDS samples ranged from 60:1-50:1 DMPC:NaK monomer.

### NMR Spectroscopy

NMR samples contained 0.5-1.4 mM NaK solubilized in q = 0.33 DMPC/DHPC bicelles in buffer containing 50-100 mM MOPS, pH 7 and 100 mM K^+^ or 600 mM Na^+^ and either 5-10% D_2_O for amide-based experiments or 100% D_2_O for methyl experiments. Samples in Na^+^ were loaded into salt-tolerant Shigemi tubes designed for high salt samples (Shigemi). Our published^25 1^H-^15^N assignments of NaK in 100 mM K^+^ at 60ºC were transferred to 40ºC for the K^+^-bound sample using an NMR temperature titration. Assignments for the K^+^-bound sample at 40ºC were then transferred to the Na^+^-bound sample and confirmed using TROSY-based HNCO and HNCA backbone-walk experiments. TROSY-based HSQC, HNCA, and HNCO spectra were collected at 40ºC on Bruker Avance III NMR spectrometers operating at 750 and 600 MHz (^1^H) and equipped with cryogenic probes. Three-dimensional spectra were acquired with non-uniform sampling (NUS) using 30-35% sampling rates and reconstructed using the SMILE algorithm implemented in nmrPipe.^50 1^H-^13^C HMQC spectra were obtained on an 800 MHz (^1^H) Varian VNMRS spectrometer equipped with a cryogenic probe. All spectra are referenced to DSS. Amide and methyl chemical shift perturbations (CSPs) were calculated using the formula, CSP = [Δδ_H_^2^ + (*W*\*Δδ_N/C_)^2^]^0.5^ where Δδ is the difference in chemical shift for a given residue between K^+^ and Na^+^ preferred states and is *W* a chemical shift weighting factor (*W*_N_ = |γ_N_/γ_H_| = 0.101, *W*_C_ = |γ_C_/γ_H_| = 0.251). Values of Δδ for Cα and C’ are presented unweighted. Note that amide CSPs were calculated on SDS back-exchanged spectra, while backbone C’ and Cα Δδ were calculated from 3D spectra obtained without SDS back-exchange, resulting in lower sequence coverage.

TROSY ^15^N R_1_, R_1_ρ, and {^1^H}-^15^N NOE experiments^28^ were recorded at 40ºC (calibrated using ethylene glycol) on a 750 MHz Bruker Avance III HD spectrometer equipped with a cryogenic probe. The number of scans was adjusted for each experiment in order to obtain satisfactory signal to noise. R_1_ and R_1_ρ values were measured using 5 relaxation delays. R_1_ experiments on the Na^+^ sample were recorded using a recycle delay of 2 s and relaxation delays of 0, 0.4,1.4, 2.4, and 3.6 s. For the K^+^ sample, the recycle delay set to 2.5 s and relaxation delays were 0.08, 0.4, 1.2, 2, and 4s. For R_1_ρ experiments on both samples the recycle delay was set to 2.5 s and relaxation delays were set to 2, 8, 12, 18, and 25 ms. The spin-lock field strength was set to 1923 Hz in both cases. The {^1^H}-^15^N NOE experiment utilized a 15s proton saturation time for the NOE experiment with an equivalent recovery delay in the reference experiment, each preceded by an additional recovery time of 1 s. The total experimental time for R_1_ and R_1_ρ experiments was approximately 3 days each, while the NOE experiment took ∼7 days.

All spectra were processed using nmrPipe^51^ and visualized using ccpNmr analysis^52^. Peak heights and the standard deviation of the baseplane noise (taken to be the error in the peak height) were extracted using ccpNmr analysis. If the error in peak height was less than 2% based on the signal to noise, an error of 2% was assumed. R_1_ and R_1_ρ values were determined by fitting the data to a single exponential decay using home-written Python scripts utilizing the *scipy*.*optimize*.*curve_fit* function. Errors were determined using 500 Monte Carlo simulations. R_1_ρ values were converted to R_2_ using the formula R_2_ = R_1_ρ/sin^2^θ -R_1_/tan^2^θ, where θ = tan- ^1^(ω/Ω), ω is the spin-lock field strength, and Ω the offset from the ^15^N carrier frequency. {^1^H}-^15^N NOE values were calculated as the ratio of peak heights in experiments with and without ^1^H saturation. The error in the NOE was calculated using standard error propagation.

3D CCH HMCQ-NOESY-HMQC spectra^53^ were obtained for samples in K^+^ and Na^+^ on an 800 MHz Varian spectrometer. Both spectra were collected with 64 (8ms) x 96 (11ms) x 1024 (^13^C-^13^C-^1^H) complex points sampled at a 35% NUS rate for Na^+^ and 40% for K^+^ and reconstructed using SMILE.^50^ The NOE mixing time was set to 200 ms and the recycle delay was set to 1 s for both samples. Total experimental time for both samples was ∼7 days.

CPMG experiments were obtained on ILV labeled NaK samples in 100 mM K^+^ and 600 mM Na^+^ purified as described above. All experiments were collected at 40ºC (calibrated using ethylene glycol). ^13^C MQ and ^1^H SQ experiments were performed using pulse sequences reported by Kay and colleagues.^34,35^ For the K^+^ sample, a ^13^C MQ dataset was recorded on 800 MHz Varian spectrometer using T_relax_ = 20 ms and ν_CPMG_ values of 50, 100, 200 (x2), 250, 300, 350, 400, 500, 600 (x2), 800, and 1000 Hz. For the Na^+^ sample, ^13^C MQ datasets were recorded on 800 and 600 MHz Varian spectrometers using T_relax_ = 30 ms and ν_CPMG_ values of 33.33, 66.67, 100, 133.33 (x2), 166.67, 200, 233.33, 266.67 (x2), 300, 333.33, 366.67, 400, 466.67, 533.33 (x2), 600, 666.67, 800, 900, and 1000 Hz. ^1^H SQ datasets for the Na^+^ sample were collected on Bruker spectrometers operating at 900 and 600 MHz using T_relax_ = 20 ms and ν_CPMG_ values of 50, 100 (x2), 150, 200, 250, 300, 350, 400 (x2), 450, 500, 600 (x2), 700, 800, 900, and 1000 Hz. The error in R_2,eff_ values was determined from replicated data points. R_ex_ values were calculated as I_(υCPMG = 50 Hz)_ – I_(υCPMG = 1000 Hz)_ for K^+^ and I_(υCPMG = 33.33 Hz)_ – I_(υCPMG = 1000 Hz)_ for Na^+^. The error in R_ex_ was calculated using standard error propagation. The Na^+ 13^C MQ and ^1^H SQ dispersion curves were fit using ChemEx.^35,54^

### Assignment of methyl resonances

In total, four separate samples were used to assign ILV methyl groups. One sample utilized the full-length NaK channel (FL-NaK) while the other three utilized NaKΔ18. These samples were expressed and purified as described above. The FL-NaK sample was produced as described above for ILV labeled samples, except 2-Keto-3-(methyl-d_3_)-butyric acid-1,2,3,4-^13^C_4_, 3-d was used as a precursor to label valines and leucines. Three 3D COSY^55^ experiments were recorded (1024×64×48 points, 8 scans, 47% non-uniform sampling), each using either one, two or three transfer steps, on an 800 MHz Varian spectrometer. The experiment allowed us to obtain correlations between valine Cγ, Cβ and Cα, as well as leucine Cδ and Cβ resonances. However, we could not obtain any Leu Cα resonances, presumably due to R_2_ relaxation during the 3-step transfer. All subsequent methyl assignment experiments were performed using NaK constructs lacking the M0 helix.

We have further produced a sample where NaK was expressed in D_2_O with ^13^C, ^1^H glucose as a carbon source and recorded 3D C-TOCSY-CHD2-HSQC experiment, as previously described^56^ (512×64×34 points, 32 scans, 47% non-uniform sampling). This experiment revealed correlations between all isoleucine, valine and leucine aliphatic carbon resonances.

Both of these experiments allowed us to determine which valine and leucine pro-S and pro-R methyl resonances belong to the same residue, as well as to distinguish valine and leucine cross-peaks in the ^1^H-^13^C HMQC spectrum. Furthermore, comparison of COSY and TOCSY experiments with data from HNCACB and HNCA experiments allowed us to correlate leucine and isolucine Cδ and valine Cγ resonances to HN, and thus determine the identity of the residue giving the origin to particular methyl group cross-peaks. Using these two experiments we could unambiguously assign all isoleucine, 50% of valine and 20% of leucine resonances. We could not unambiguously assign the remaining residues either due to the lack of resolution in our 3D spectra, or the lack of Cβ cross-peaks in HNCACB experiment.

To obtain the full assignment, we have recorded 3D CCH HMQC-NOESY-HMQC experiment displaying correlations between spatially proximal methyl groups, as described above, and used MAGMA^57^ without using the restraints from COSY and TOCSY experiments, except differentiating leucine and valine cross-peaks. The 3E8H NaK crystal structure was used to generate the connectivity graph. MAGMA produced confident assignments for 20 out of 30 assignable residues. Two residues, L48 and V45 are too far away from any other methyl group, and thus were not included in the calculation. Comparison of data obtained with COSY and TOCSY experiments were in an excellent agreement, and remaining ambiguous assignments proposed by MAGMA could be easily resolved.

Lastly, we performed a 3D ^1^H-NOESY-^1^H,^15^N-HSQC experiment on the same sample as was used to record 3D CCH HMQC-NOESY-HMQC experiment to validate our assignment. The experiment was performed with 46% non-uniform sampling, with 1024×96×40 points, 32 scans, 140ms mixing time. This experiment reveals correlations between nearby methyl and amide groups. The experiment was in complete agreement with our methyl assignment data.

## ACKNOWLEDGEMENTS

We would like to thank Tairan Yuwen for providing the ChemEx module for fitting ^1^H CPMG data and for useful discussions on performing the fitting. Research reported in this publication was supported by NIGMS of the National Institutes of Health under award numbers R01GM116047 and R35GM141748. The content is solely the responsibility of the authors and does not necessarily represent the official views of the National Institutes of Health. This study made use of the National Magnetic Resonance Facility at Madison, which is supported by NIH grants P41GM103399 (NIGMS) and P41GM66326 (NIGMS). Additional equipment was purchased with funds from the University of Wisconsin, the NIH (RR02781, RR08438), the NSF (DMB-8415048, OIA-9977486, BIR-9214394, and the USDA.

## AUTHOR CONTRIBUTIONS

A.L. prepared NaK samples, collected and analyzed NMR data, and wrote the original manuscript. V.K. prepared NaK samples, performed methyl group assignments, and revised the manuscript. M.T. assisted in NMR data collection and revised the manuscript. K.H.W. supervised the project, acquired funding, and revised the manuscript.

