## Supplementary Information for "Ion-dependent structure, dynamics, and allosteric coupling in a non-selective cation channel"

*Supplementary note 1: SDS-mediated back-exchange enables solution NMR studies of NaK*

As noted in the text, SDS-mediated refolding and back-exchange greatly increased the number of peaks visible in perdeuterated NaK samples (Fig. S1). For $K^+$, most residues are at least partially back exchanged, enabling assignment of the majority of peaks in SDS back-exchanged spectra. Assignments were also aided by previous 3D experiments collected at 60ºC on non-deuterated samples where more signals are present, albeit with lower s/n than for deuterated samples. However there are four peaks that appear that remain unassigned, as 3D experiments were not performed on back-exchanged samples. It is likely that these peaks correspond to the four unassigned residues, 57, 60, 61, and 63. Notably, the peak observed for T62 and one of the new peaks that appear are quite weak, suggesting that there could be intermediate timescale dynamics occurring in this region or that H/D exchange is not complete in this region even in SDS. In the case of $Na^+$, as well, we lack assignments for residues 57 and 60-63 for much the same reasons as for $K^+$. Additionally, for $Na^+$ numerous minor states are observed at low contour levels, which could be an artifact of SDS membrane extraction. We stress, though, that the major state observed in SDS back-exchanged spectra overlays very well with the spectrum obtained without SDS treatment. This issue is not observed in $K^+$.

*Supplementary note 2: $^{15}N$ relaxation studies of NaK*

$^{15}N$ $R_2$ data collected for NaK in 100 mM $K^+$ and 600 mM $Na^+$ are shown plotted against the NaK sequence in Fig. S2. It is clear from the plots in Fig. 2 and Fig. S2 that there is a marked increase in $R_1$ and decreases in the NOE and $R_2$ at the ends of the M1 (~18-23) and M2 (~100-110) helices in both salt conditions; this is commonly observed for proteins and is indicative of fraying at the ends of helices. It is also evident that, in general, the measured $R_2$ values are higher for the $Na^+$ sample. This is likely due to differences in bicelle size and/or solution viscosity arising from differences in the protein:lipid ratio used for reconstitution (1:60 for $K^+$, 1:95 for $Na^+$). Apparent values of the global rotational correlation time ($\tau_c$) were calculated for residues with NOE values >0.65 and ~average $R_2/R_1$ values, as these residues are unlikely to undergo significant internal dynamics.[1,2] Residue-specific values of $\tau_c$ were estimated from the $R_2/R_1$ ratio for qualifying residues, and then averaged to obtain a global estimate for $\tau_c$. There is a small but significant difference between the $K^+$ sample, $\tau_c = 43 \pm 2$ ns, and the $Na^+$ sample, $\tau_c = 47 \pm 2$ ns.

The trend observed in the SF for $R_1$ and NOE data is generally maintained for $R_2$, although direct comparison between $K^+$ and $Na^+$ conditions is hampered by the aforementioned offset. $R_2$ values in $K^+$ for D66 and N68 are elevated relative to the values observed for G67, G69, and S70, whereas $R_2$ values are generally consistent for residues 66-70 in $Na^+$. This could be indicative of slower internal dynamics contributing to $R_2$ for D66 and N68 in $K^+$.

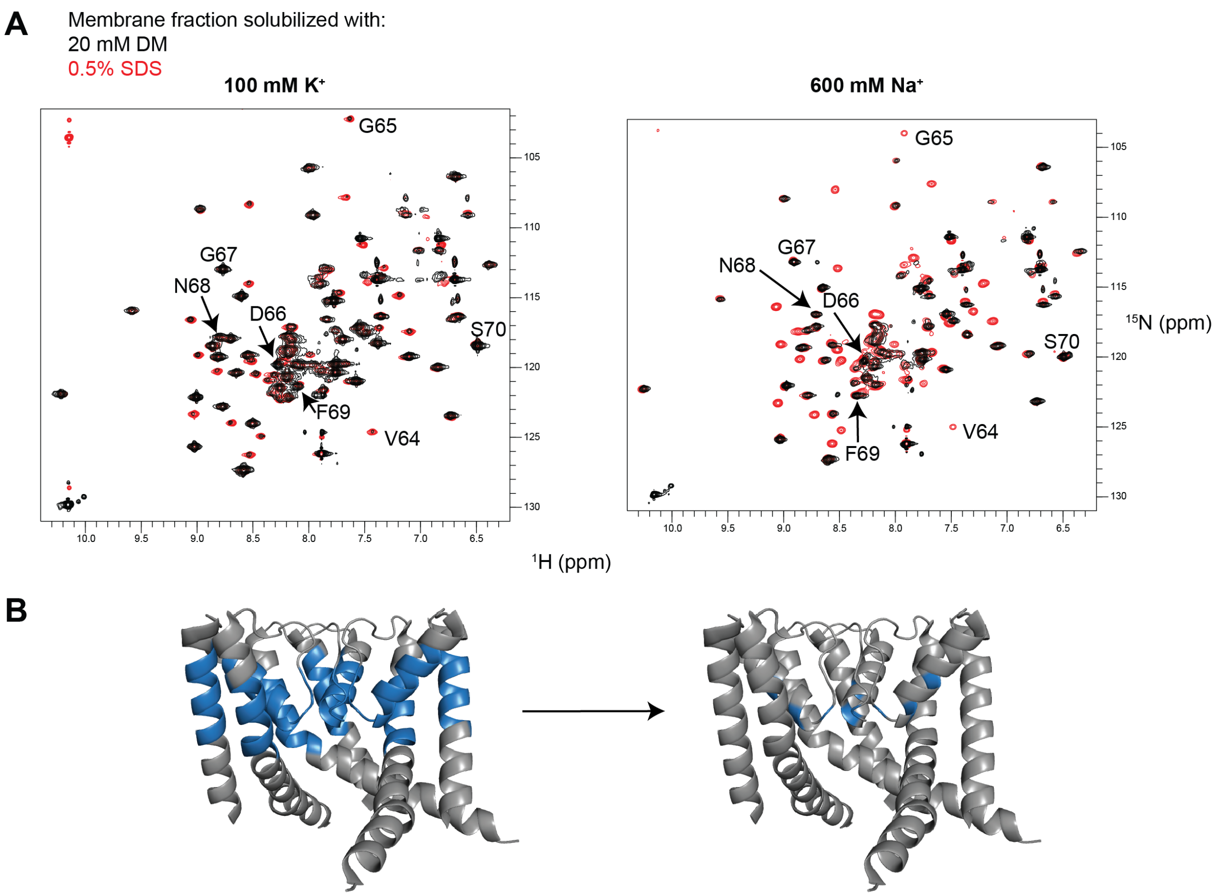


**Figure S1. Back-exchange of amide protons increases when the membrane fraction is solubilized in 0.5% SDS. a** Overlay of $^{1}H$-$^{15}N$ TROSY-HSQC spectra of NaK in 100 mM $K^+$ (left) and 600 mM $Na^+$ (right), where the membrane fraction was solubilized with 20 mM DM (black) or 0.5% SDS (red). All spectra were obtained with NaK reconsituted into DMPC:DHPC bicelles (q ~ 0.33). Residues in the SF are indicated; note that V64 and G65 are poorly back-exchanged without SDS membrane extraction. The sidechain peak for W19 is aliased in the back-exchanged spectra. **b** SDS solubilization promoted back-exchange in NaK. On the left, regions with poor back exchange are colored in blue. On the right, residues that remain without assignment after SDS solubilzation are colored blue. Residues turning from blue to gray increase in intensity after solubilization with SDS, indicating increased back-exchange.

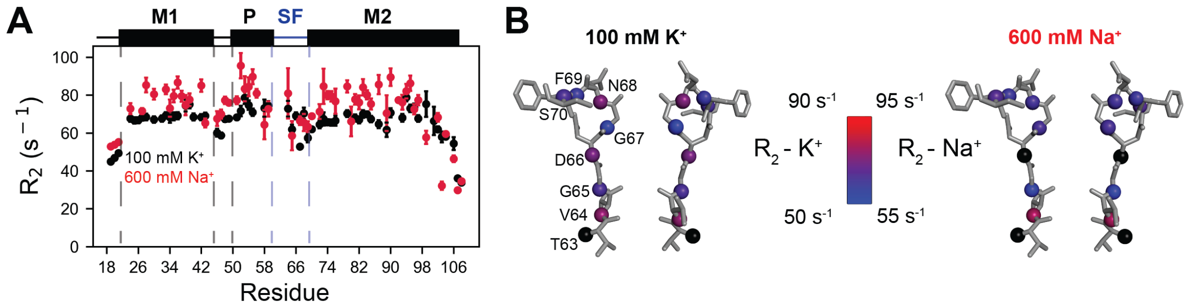


**Fig. S2 $^{15}$N $R_2$ values vary across NaK SF. a** Backbone amide $^{15}$N $R_2$ values recorded at 750 MHz for NaK in 100 mM $K^+$ (black) and 600 mM $Na^+$ (red). $R_2$ values are calculated from $R_1$ and $R_1\rho$ as described in the Methods. Secondary structure elements are shown above, where helices are rectangles and loops are lines. The boundaries between elements are based on the 3E8H crystal structure. Residues 63-70 are indicated by the blue lines and are shown as sticks in B. **b** Relaxation data plotted on the NaK SF (showing 2 of 4 opposing subunits). Backbone N atoms are shown as spheres and are colored according to the scale indicated for each parameter. Residues for which data are unavailable due to resonance overlap or lack of assignment are colored black.

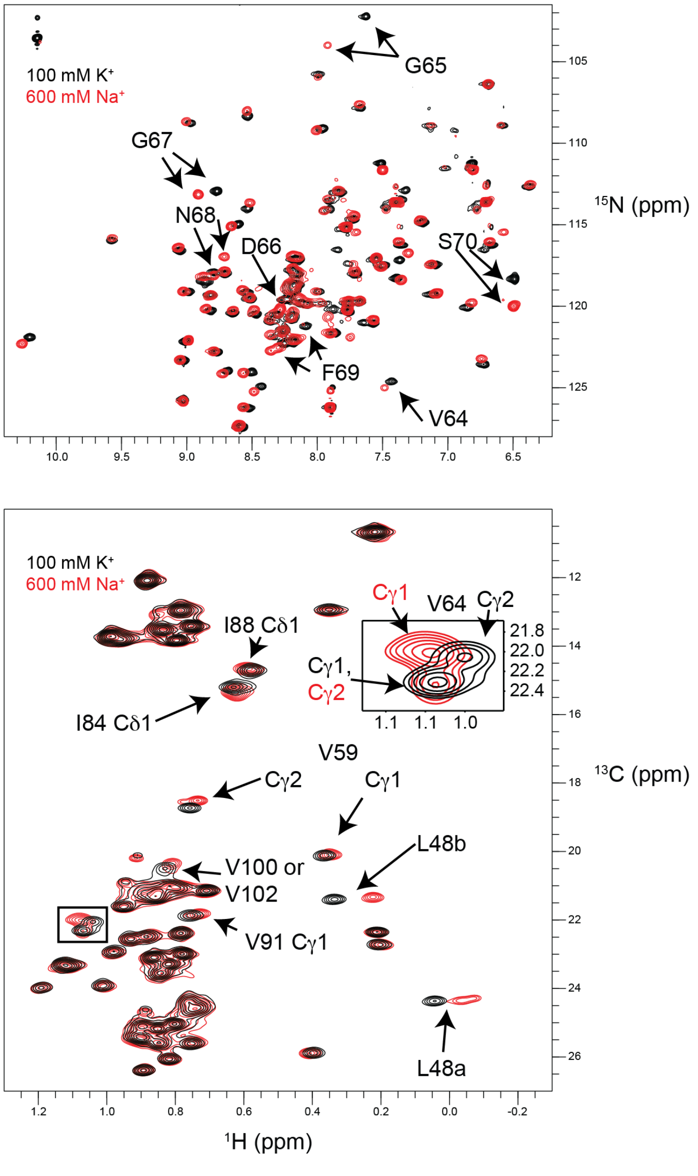


**Figure S3. Overlay of NaK spectra in 100 mM $K^+$ vs. 600 mM $Na^+$. a** TROSY-HSQC spectra of NaK in the indicated ionic condition. Residues in the SF that undergo significant chemical shift perturbations are indicated. Note that the W19 sidechain amide peak is aliased in these spectra. **b** $^1H$-$^{13}C$ HMQC spectra of ILV-labeled NaK in the indicated ionic conditions. Peaks near the SF that shift between conditions are labeled. Methyl resonances that are stereospecifically assigned are labeled according to the PDB labeling for 3E8H, while resonances without stereospecfic assignments are labeled a or b. Resonances for V64 are expanded in the inset and labels are color-coded red for $Na^+$ and black for $K^+$.

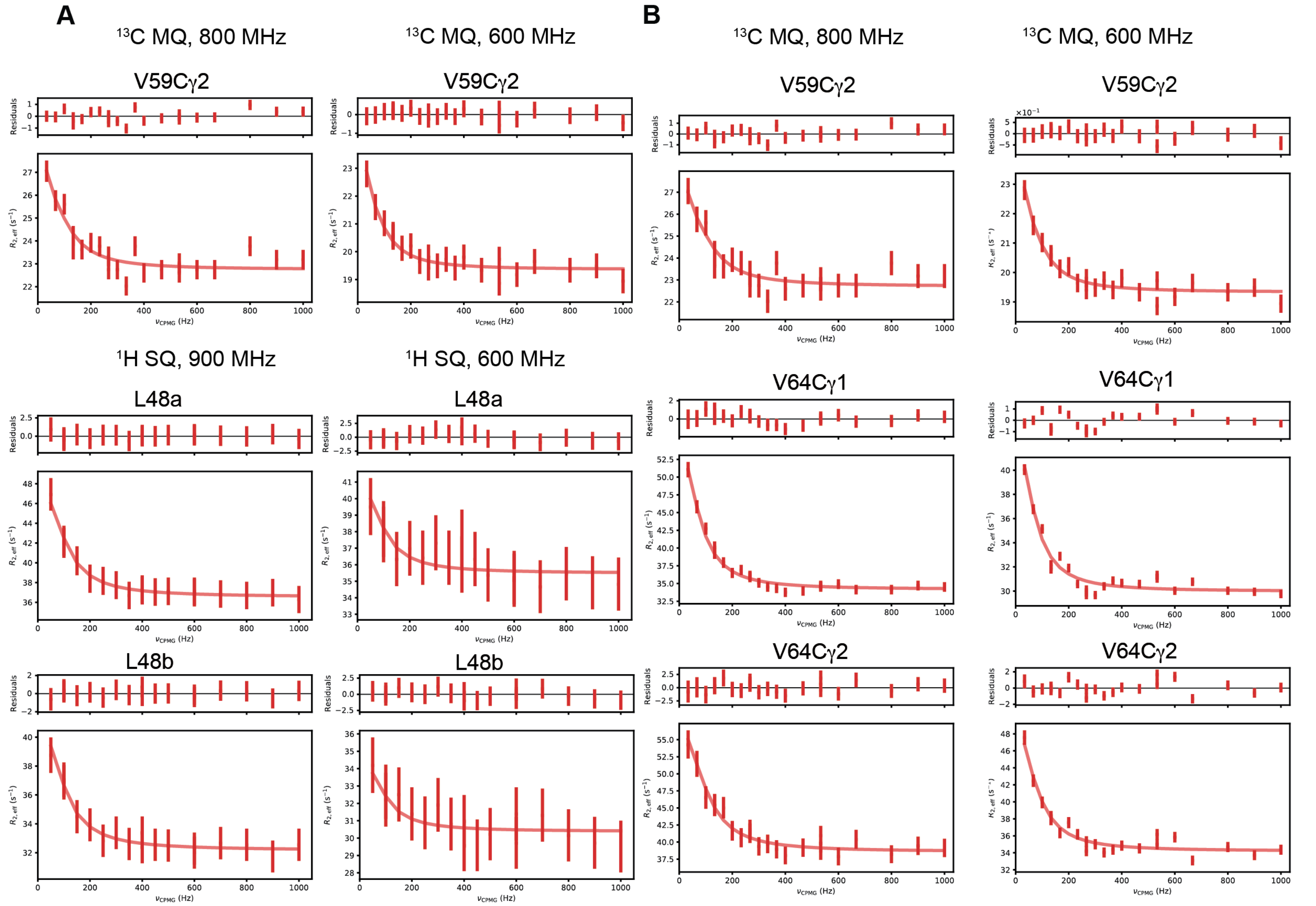


**Figure S4. Fit results for global fitting of CPMG data. a** Fit results for simultaneous fitting of V59 $^{13}$C MQ and L48 $^{1}$H SQ CPMG data. The extracted value of $k_{ex}$ = 356 ± 67 $s^{-1}$. **b** Fit results for simultaneous fitting of $^{13}$C MQ data for V59 and V64. The extracted value of $k_{ex}$ = 390 ± 47 $s^{-1}$.
